# Molecular classification reveals the diverse genetic features and prognosis of gastric cancer: a multi-omics consensus ensemble clustering

**DOI:** 10.1101/2021.06.07.447364

**Authors:** Xianyu Hu, Zhenglin Wang, Qing Wang, Ke Chen, Qijun Han, Suwen Bai, Juan Du, Wei Chen

## Abstract

**Background:** Gastric cancer (GC) is the fifth most common tumor around the world, it is necessary to reveal novel molecular subtypes to guide the selection of patients who may benefit from specific target therapy.

**Methods:** Multi-omics data, including RNA-sequence of transcriptomics (mRNA, LncRNA, miRNA), DNA methylation and gene mutation of TCGA-STAD cohort was used for the clustering. Ten classical clustering algorithms were applied to recognize patients with different molecular features via the R package “*MOVICS*”. The activated signaling pathways were evaluated using the single-sample gene set enrichment analysis. The difference distribution of gene mutations, copy number alterations and tumor mutation burden was compared, and potential response to immunotherapy and chemotherapy was assessed as well.

**Results:** Two molecular subtypes (CS1 and CS2) were recognized by ten clustering algorithms with further consensus ensembles. Patients in the CS1 group were found to contain a shorter average overall survival time (28.5 vs. 68.9 months, *P* = 0.016), and progression-free survival (19.0 vs. 63.9 months, P = 0.008) compared to the CS2 group. CS1 group contained more activation of extracellular associated biological process, while CS2 group displayed the activation of cell cycle associated pathways. The significantly higher total mutation numbers and neo antigens were observed in CS2 group, along with the specific mutation of TTN, MUC16 and ARID1A. Higher infiltration of immunocytes were also observed in CS2 group, reflected to the potential benefit from immunotherapy. Moreover, CS2 group also can response to 5-fluorouracil, cisplatin, and paclitaxel. The similar diverse of clinical outcome of CS1 and CS2 groups were successfully validation in external cohorts of GSE62254, GSE26253, GSE15459, and GSE84437.

**Conclusion:** Novel insight into the GC subtypes was obtained via integrative analysis of five omics data by ten clustering algorithms, which can provide the idea to the clinical target therapy based on the specific molecular features.

## Introduction

Gastric cancer (GC) is the fifth most common tumor and ranks third in mortality worldwide[1]. China is the most affected country by GC, accounting for 42.6% of global incidence and 45% of GC-related death. In addition, it is second in terms of incidence among all malignancies in China[2, 3]. Helicobacter pylori infection is the most important pathogenic factor of GC. Most patients infected with Helicobacter pylori remain asymptomatic, however, almost all patients’ symptoms are accompanied with varying degrees of atrophic gastritis and intestinal metaplasia[4]. For most early GC patients, endoscopic resection is a definitive treatment[5]. For non-early GC patients, gastrectomy with D2 lymphadenectomy and chemotherapy is an internationally agreed upon treatment[6–8]. Recently, combination adjuvant chemotherapy in both the preoperative and postoperative period have been considered in being beneficial for GC patients[9, 10].

The Cancer Genome Atlas (TCGA) group has divided four molecular subtypes of GC, including Epstein-Barr virus (EBV)-positive, microsatellite instability (MSI), genomically stable (GS) and chromosomal instability (CIN)[11]. The EBV-positive subtype has demonstrated enrichment of 9p amplification with consecutive high expression of PD-L1 and PD-L2. PD-1/PD-L1 inhibitors, which may have alleviating effects in EBV-positive GC[12]. The MSI subtype is characterized by mutations in the DNA mismatch repair system (MMR) or silencing of respective promoter regions by hypermethylation, which can detect and repair DNA base pair mismatches. In patients with resectable primary GC, MSI serves as a robust biomarker that indicates favourable post-surgical survival outcomes[13]. GS is generally a diffuse GC, in which CDH1 and RHOA mutations are detectable or frequently show CLDN18-ARHGAP fusion[14]. Moreover, TP53 mutations are found in 70% of CIN GCs, which may result in aneuploidy and focused amplification of receptor tyrosine kinases[15]. The Asian Cancer Research Group (ACRG) has reported four expression subtypes of GC: MSS/TP53-, MSS/TP53+, MSI, and MSS/EMT, in which MSS refers to microsatellite stable tumors. Among these molecular subtypes, MSI has the best prognosis, while MSS/EMT has the poorest prognosis and MSS/TP53- and MSS/TP53+ have intermediate prognosis[16].

Currently, only two biomarkers are available to predict the therapeutic effect of patients: human epidermal growth factor receptor 2 (HER2) for trastuzumab and programmed death-ligand 1 (PD-L1) expression for pembrolizumab[17–19]. Although several immunotherapy-based trials have reported widespread tumor response rates in GC patients, many phase III trials of targeted drugs have not shown significant survival benefits[20, 21]. Weak selection of patient molecules included in clinical trials remains an issue and may limit the evaluation of the benefits of many therapeutic drugs, such as anti-angiogenic molecules and newer immunomodulatory agents. Therefore, it is necessary to uncover novel biomarkers in order to better select patients who may benefit from specific target therapy.

## Methods

### Preparing the multi-omics data of the TCGA-STAD cohort

The TCGA-STAD cohort was chosen as the discovery cohort as it has multi-omics data, such as the bulk RNA-sequence of transcriptomics (mRNA, LncRNA, miRNA), DNA methylation and gene mutation. In addition, the expression data of transcriptomics were downloaded using the R package “TCGAbiolinks”, while the gene symbols of mRNA were annotated using the GENCODE27 annotation file and the LncRNAs were identified by Vega. The expression of miRNAs was downloaded from UCSC Xena (), in which the mature miRNA names were renamed using the R package “miRNAmeConverter” along with the miRbase 21.0. The counts data of transcriptomics were then transferred to the transcripts per kilobase million (TPM) value for subsequent analysis. Additionally, Illumina DNA methylation 450 data was downloaded from UCSC Xena, while somatic mutations were obtained from the cBioPortal (). The clinicopathological information of the TCGA-STAD cohort was also obtained using “TCGAbiolinks”. A total of 243 GC patients were finally enrolled, who all possessed the types of omics data.

### Preparing the external validation cohort

In order to verify the new finding from the discovery cohort, the mRNA expression matrix and clinical information were also downloaded from four external GEO cohorts, including GSE62254, GSE26253, GSE15459 and GSE84437. The GSE62254 cohort contained the microarray profiles from 300 histologically confirmed gastric adenocarcinoma patients from the ACRG gastric cohort. Meanwhile, the GSE26253 cohort contained a microarray gene expression profile of archived paraffin-embedded tumor blocks from 432 gastric adenocarcinoma patients. GSE15459 recorded a genome-wide mRNA expression profile of 192 primary gastric tumors from Singapore patients. Moreover, GSE84437 contained the data of 433 gastric tumor patients from Yonsei University. The microarray platform of GSE62254 and GSE26253 was obtained from Affymetrix Human Genome U133 Plus 2.0 Array, Illumina HumanRef-8 WG-DASL v3.0 for GSE15459, and Illumina HumanHT-12 V3.0 expression beadchip for GSE84437.

### Identifying molecular subtypes via integrative analysis

In conjunction with guidelines from the recently published R package “*MOVICS*”[22], the molecular subtypes were identified according to the above-mentioned multi-omics data. The OS related factors were first evaluated by univariate Cox regression, including mRNA, LncRNA, miRNA and DNA methylation CpG sites. In regard to the mutant genes, genes with a mutant frequency higher than 10% were selected. In order to find the appropriate number of subtypes, analysis of clustering prediction index (CPI) [23] and Gaps-statistics[24] based on the multi-omics data were performed. Subsequently, ten clustering algorithms were used to separate the patients according to different subtypes, after which the combined classification was finally obtained in view of the concept of consensus ensembles to identify the subtypes with high robustness. The ten clustering algorithms contained iClusterBayes, moCluster, CIMLR, IntNMF, ConsensusClustering, COCA, NEMO, PINSPlus, SNF, and LRA. Quantification of sample similarity in the subtypes was also assessed using the silhoutte score.

### Evaluating the activation of signaling pathways and immune infiltration

The subtype-specific upregulated and downregulated genes were calculated using the R package “limma”, with significant threshold in both the nominal and adjusted P value < 0.05. The activation of Gene Ontology (GO) terms of each patient were then evaluated using the single-sample gene set enrichment analysis (ssGSEA) by the R package “GSVA”. The primary result of the gene set enrichment analysis is the enrichment score (ES), which reflects the degree to which a gene set is overrepresented at the top or bottom of a ranked list of genes. A positive ES indicates gene set enrichment at the top of the ranked list, while a negative ES indicates gene set enrichment at the bottom of the ranked list. The normalized enrichment score (NES) is the primary statistic for examining gene set enrichment results. The false discovery rate (FDR) is the estimated probability that a gene set with a given NES represents a false positive finding. The nominal p value estimates the statistical significance of the enrichment score for a single gene set. The gene sets of the GO biological processes were downloaded from the Molecular Signatures Database (MSigDB, https://www.gsea-msigdb.org/gsea/msigdb/index.jsp). The specific pathways of one subtype should be significantly different from the other subtypes. The mean of the enrichment score for each subtype was used in the visualization of the heatmap. The gene sets of immune signatures and stromal signatures were obtained from a previous study[25], and the NES score was calculated in order to compare the different immune activated statuses among the different subtypes.

### Evaluating the response to immunotherapy and chemotherapy

In order to evaluate the individual likelihood of responding to immunotherapy, the Tumor Immune Dysfunction and Exclusion (TIDE) algorithm was employed. Moreover, according to a melanoma cohort who received anti-CTLA-4 or anti-PD-1 checkpoint inhibition therapy, specific gene sets were obtained with 795 genes[26]. Subclass mapping analysis was then conducted in order to compare the similarity of risk groups with the immunotherapy subgroups and point out the responders of anti-CTLA-4 or anti-PD-1 immunotherapy[27]. The chemotherapeutic response for each sample was then predicted based on the Genomics of Drug Sensitivity in Cancer (GDSC). Accordingly, 3 commonly used chemo drugs, 5-Fluorouracil, cisplatin, and paclitaxel were selected. The response to the above six drugs was compared by estimating the samples’ half-maximal inhibitory concentration (IC_50_) via ridge regression.

### Characteristics of genetic alterations among subtypes

The number of nonsynonymous mutations per million bases was then calculated to evaluate the tumor mutation burden (TMB), while FGA referred to the percentage of genome affected by copy number gains or losses. The total mutation numbers and neo antigens were obtained from a previous study[28]. Oncoprint mutation landscape was generated using the R package “maftools” [29]. The cumulative recurrent events were then displayed using the R package “*Survival*”. The copy number segment data were downloaded from FireBrowse (http://firebrowse.org/) and displayed by the R package “maftools”.

### Statistical analysis

All analyses were completed using the R software v4.0.3 (http://www.r-project.org). Comparisons of continuous data between two groups were performed using the Student’s T-test or Wilcoxon test. Pearson correlation coefficient test was then employed in order to assess the relationship between the two factors. The distribution of categorical variables between groups were then compared using the Chi-square test. K-M survival analysis was performed to explore the survival difference between high-risk score group and low-risk score. A log-rank test was used to estimate the survival analysis. HR and 95%CI were calculated by the cox model. Meanwhile, the independent prognostic effect of risk score was calculated by multivariate cox regression analysis. P < 0.05 was considered to be a statistically significant difference.

## Results

### GC patients were separated to two subtypes by MOVICS

In view of the results of CPI and Gaps-statistics analysis, the highest average value of these two methods when the number of subtypes was found to be 2 (Figure 1A). Subsequently, the integration of the clustering results derived from the 10 algorithms was performed via consensus ensembles. To make the classification more robust (Figure 1B), patients were divided into cluster 1(CS1) and cluster 2 (CS2). Moreover, the similarity of the samples in the identified clusters was also evaluated by silhouette analysis, in which the silhouette score of CS1 was found to be 0.55 while that of CS2 was 0.61. Accordingly, the results demonstrated that the clusters were well apart from each other and were clearly distinguished (**Figure 1C**). In regard to the distribution of the multi-omics data for the classification, the different distribution of mRNA, LncRNA, miRNA, DNA methylation CpG sites and mutant genes were visualized, as shown in **Figure 1D**, of which the top 10 OS-associated factors of each omics are displayed in the right, while the clinical features of grade, stage, age and gender are also listed.

**Figure 1.**
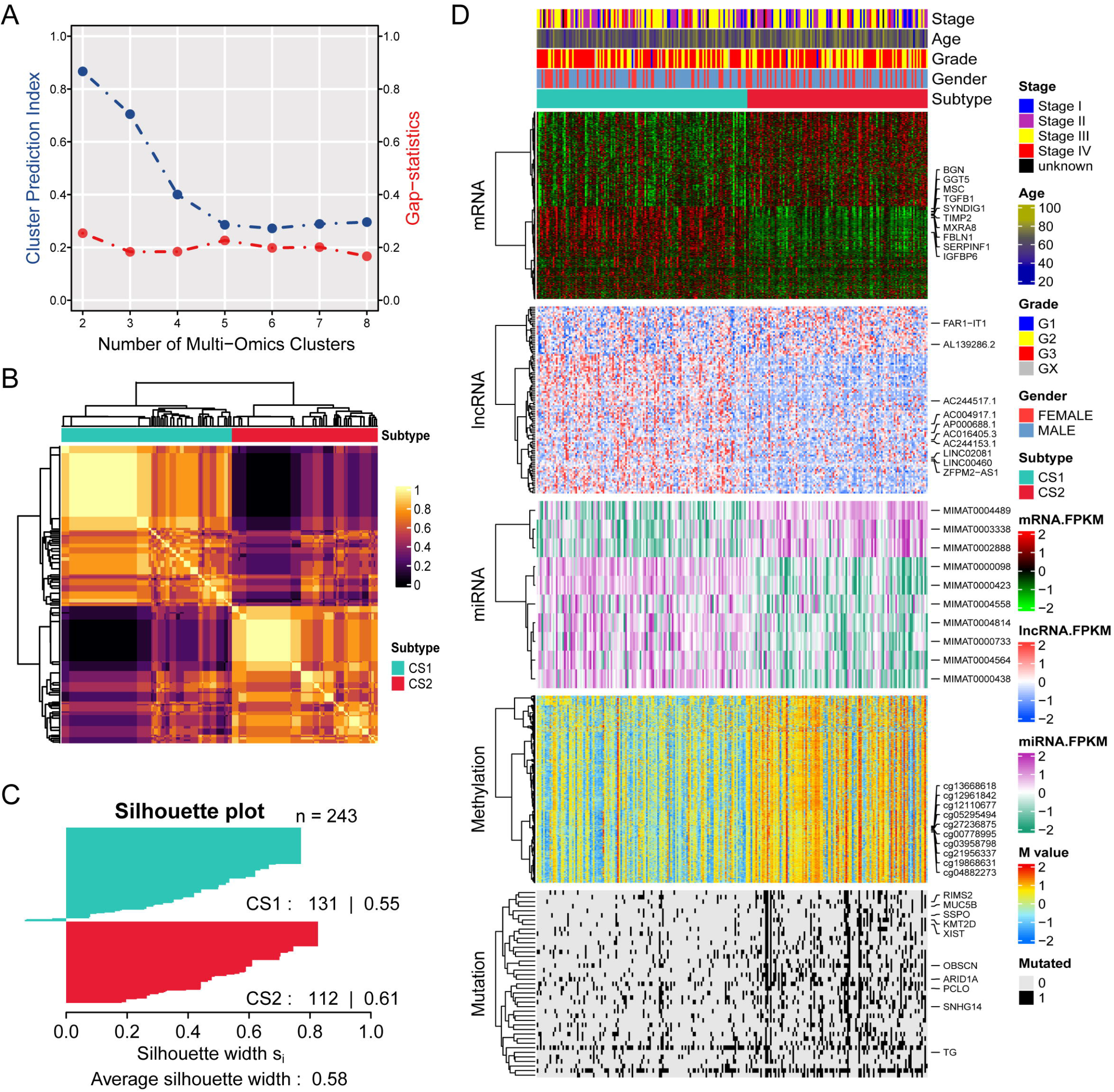
Identifying the molecular clusters. A. Prediction of optimal cluster number of multi-omics clusters by cluster prediction index and Gap-statistics; B. Consensus heatmap based on the 10 integrative clustering algorithms to refine the clusters; C. Quantification of sample similarity using silhouette score based on the consensus ensembles result; D. Comprehensive heatmap for the enrolled multi-omics data with the annotation of clinical features.

### CS1 patients suffered from shorter overall survival time and progression-free survival

The clinical prognosis outcome of GC patients in CS1 and CS2 were further compared. Accordingly, patients in the CS1 group were found to contain a shorter average OS time compared to the CS2 group (28.5 vs. 68.9 months, P = 0.016, **Figure 2A**). A similar difference in PFS time was also observed, in which the average PFS time of the CS1 group was significantly shorter than that of the CS2 group (19.0 vs. 63.9 months, P = 0.008, **Figure 2B**). The distribution of different clinicopathological features in CS1 and CS2 groups were also compared (**Table 1**). Patients in the CS1 group contained more white patients (P = 0.007), more patients live with tumors (P = 0.045), and low average age (P = 0.027), while the distribution of tumor stage, grade and gender showed no differences (all P > 0.05). The distribution of different factors in separated subtypes are given in **Figure 2C**.

**Figure 2.**
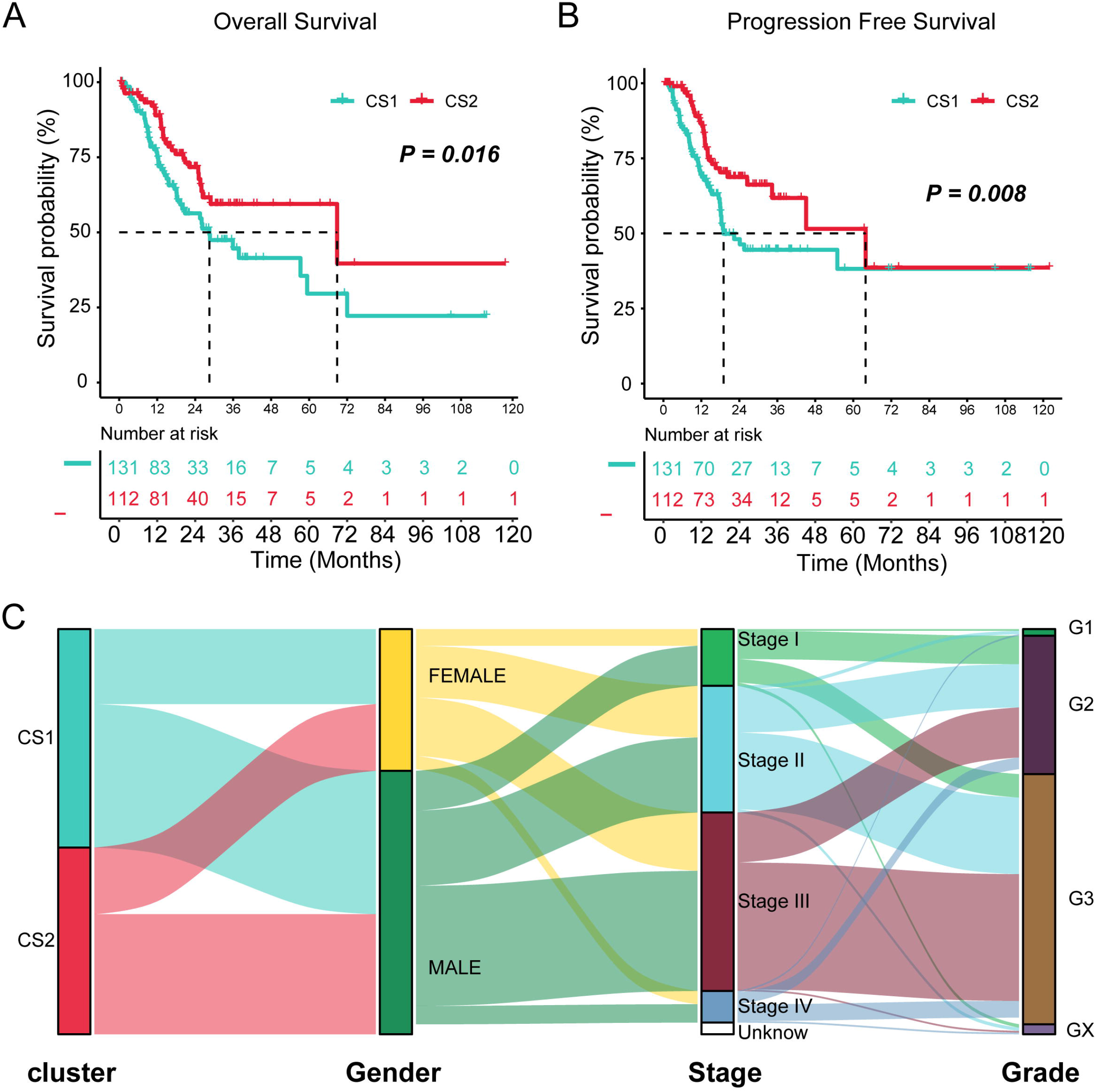
Differential clinical outcome of two groups. A. Kaplan-Meier survival analysis of the overall survival of the two groups; B. Kaplan-Meier survival analysis of the progression free survival of the two groups; C. The distribution of the clusters with different clinical features.

### Different expressed genes and signaling pathways in CS1 and CS2

To characterize the features of CS1 and CS2, the different expressed genes (DEGs) between the CS1 and CS2 groups were first identified. Based on the “limma” package, 2016 upregulated genes were selected from the CS1 group, while 400 upregulated genes were taken from the CS2 group with a threshold P value < 0.05 and fold-change > 2 (**Table S**). With the help of Metascape, the DEGs were annotated and enriched to biological processes. For CS1 activated pathways, the activation of extracellular associated biological process were mostly observed, including epithelial mesenchymal transition, cell junction organization, cellular component morphogenesis, response to growth factor and cell-substrate adhesion pathways (**Figure 3A**). In regard to the CS2 group, pathways of cell cycle, including G2M checkpoint, cell cycle, E2F targets, G1/S-specific transcription, DNA replication and repair biologic process were mainly focused on (**Figure 3B**).

**Figure 3.**
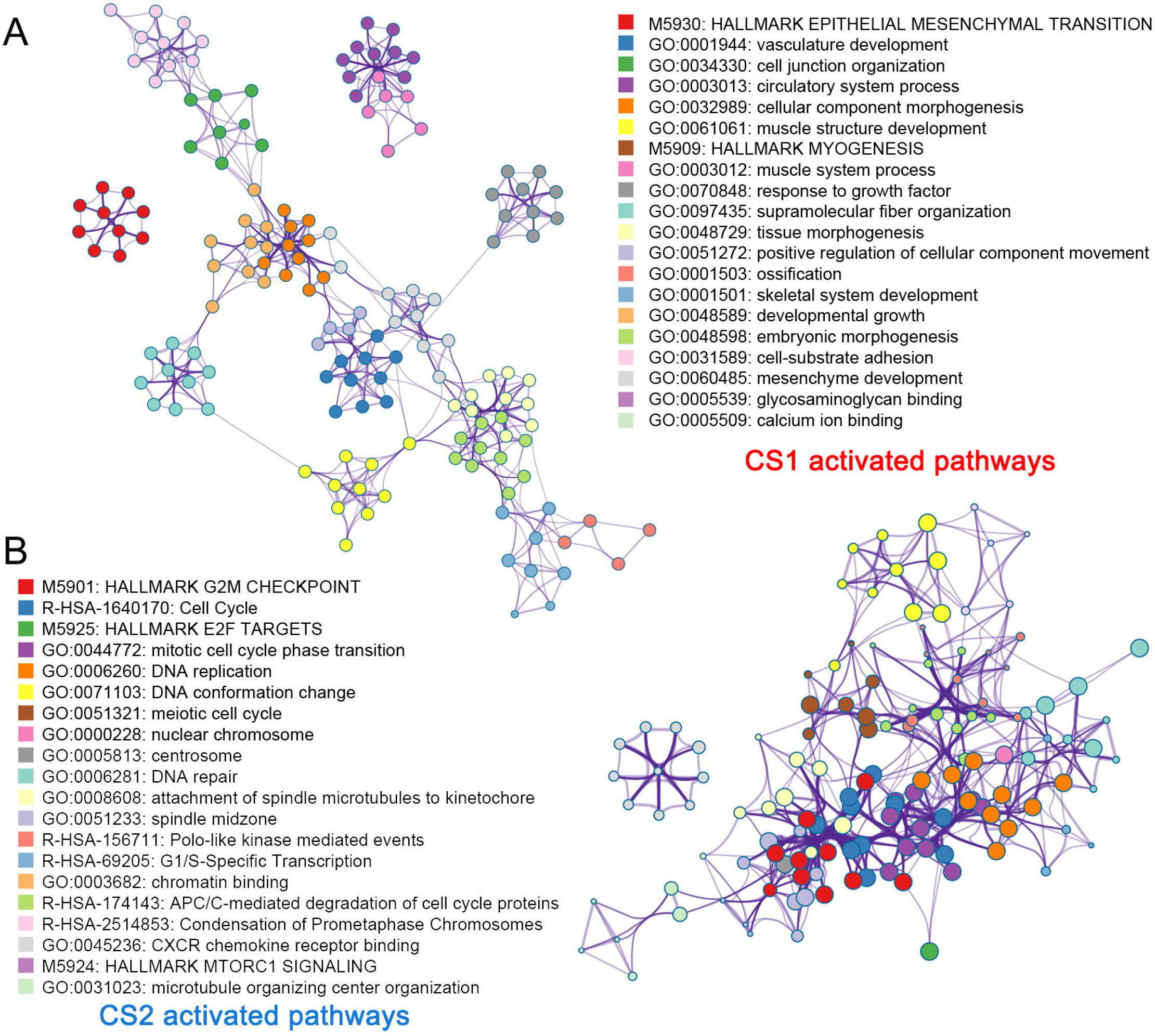
Cluster-specific biomarkers and signaling pathways. A. The activated signaling pathways in the CS1 group enriched and annotated by 2016 significantly upregulated genes; B. The activated signaling pathways in the CS2 group enriched and annotated by 2016 significantly upregulated genes.

For all of the DEGs among CS1 and CS2, this study focused on SMOC2 (**Figure 4A**). SMOC2 was found to be significantly increased in the CS1 subtype, with a fold-change of 4.537 (adjusted P value < 0.001). The different activation of extracellular matrix associated pathways were also assessed by GSEA analysis, in which three pathways were found to be significantly activated in the CS1 subtype compared to the CS2 subtype (all P < 0.01, **Figure 4B**). Several studies have reported the function of SMOC2 in tumorigenesis through extracellular matrix associated pathways, however, few were concerned with GC patients. Accordingly, a high expression of SMOC2 was observed to be linked with poor prognosis in GC patients (HR = 1.66, 95% CI = 1.081-2.551, P = 0.021, **Figure 4C**). Moreover, the mRNA expression of SMOC2 was found to be positively associated with mesenchymal maker, Vimentin (R = −0.32, P < 0.001, **Figure 4D**), which was negatively associated with epithelia marker, E-Cadherin (R = −0.27, P < 0.001, **Figure 4D**). In addition, the prognostic value of SMOC2 in three GEO cohorts were also evaluated, in which high levels of mRNA of SMOC2 was found to be associated with poor prognosis (GSE15459, HR = 1.56, 95% CI = 1.05-2.32, P = 0.027; GSE62254, HR = 2.02, 95% CI = 1.40-2.91, P < 0.001; GSE51105, HR = 1.52, 95% CI = 0.91-2.54, P = 0.11; **Figure 4E**).

**Figure 4.**
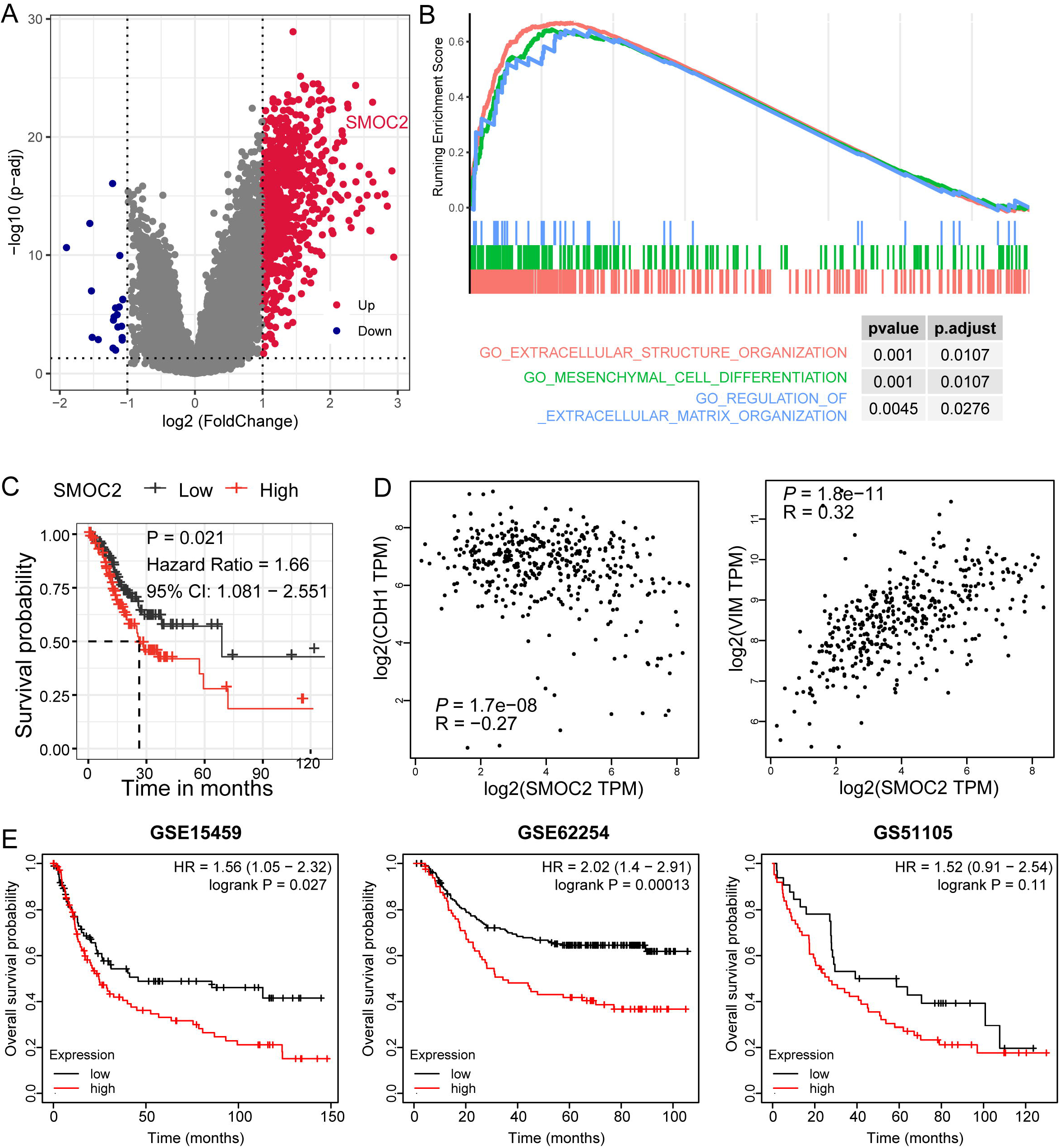
SMOC2 and extracellular pathways activated in CS2 cluster. A. Differently expressed genes among CS2 to CS1 groups; B. Activation of extracellular associated signaling pathways; C. High level of SMOC2 indicated poor prognosis in the TCGA-STAD cohort; D. The expression of SMOC2 positively correlated with epithelial mesenchymal transition biomarkers; E. High level of SMOC2 indicated poor prognosis in the GSE15459, GSE62254, GSE51105 cohorts.

### CS2 contained more genetic alterations

The gene mutation and copy number alteration play pivotal role in the initial stages and development of tumorigenesis. In this regard, the genetic alterations among CS1 and CS2 were then compared.

The number of nonsynonymous mutations per million bases was calculated so as to evaluate the TMB. Here, a significantly higher average TMB value was present in CS2 compared to that of CS1 (log10 transformed TMB value: 0.68 vs. 0.52, P < 0.001, **Figure 5A**). Moreover, the total mutation numbers in CS2 were also higher than that of the CS1 group (P < 0.001, **Figure 5B**), in conjunction with the numbers of neo antigens (P < 0.001, **Figure 5C**). Additionally, the difference of mutation in a single gene was also compared, where there were more TTN mutations (65.2% vs. 42.7%, P < 0.001), MUC16 mutations (44.1% vs. 22.1%, P = 0.002), and ARID1A mutations (36.6% vs. 11.5%, P < 0.001) (**Figure 5D, Table 2**). Mutated MUC16 decreased the cumulative rate of OS events, which acted as a tumour suppressor (P = 0.012, **Figure 5E**). Based on the databases from cBioPortal, the total genetic alteration of MUC16 was found to be about 15% to 30%, in which the altered group demonstrated better prognosis (P < 0.001, **Figure S1**). The copy number alterations were also calculated, in which the CS2 groups were noted to contain more CNAs of both genomes lost or gained (all P < 0.05, **Figure 5F**).

**Figure 5.**
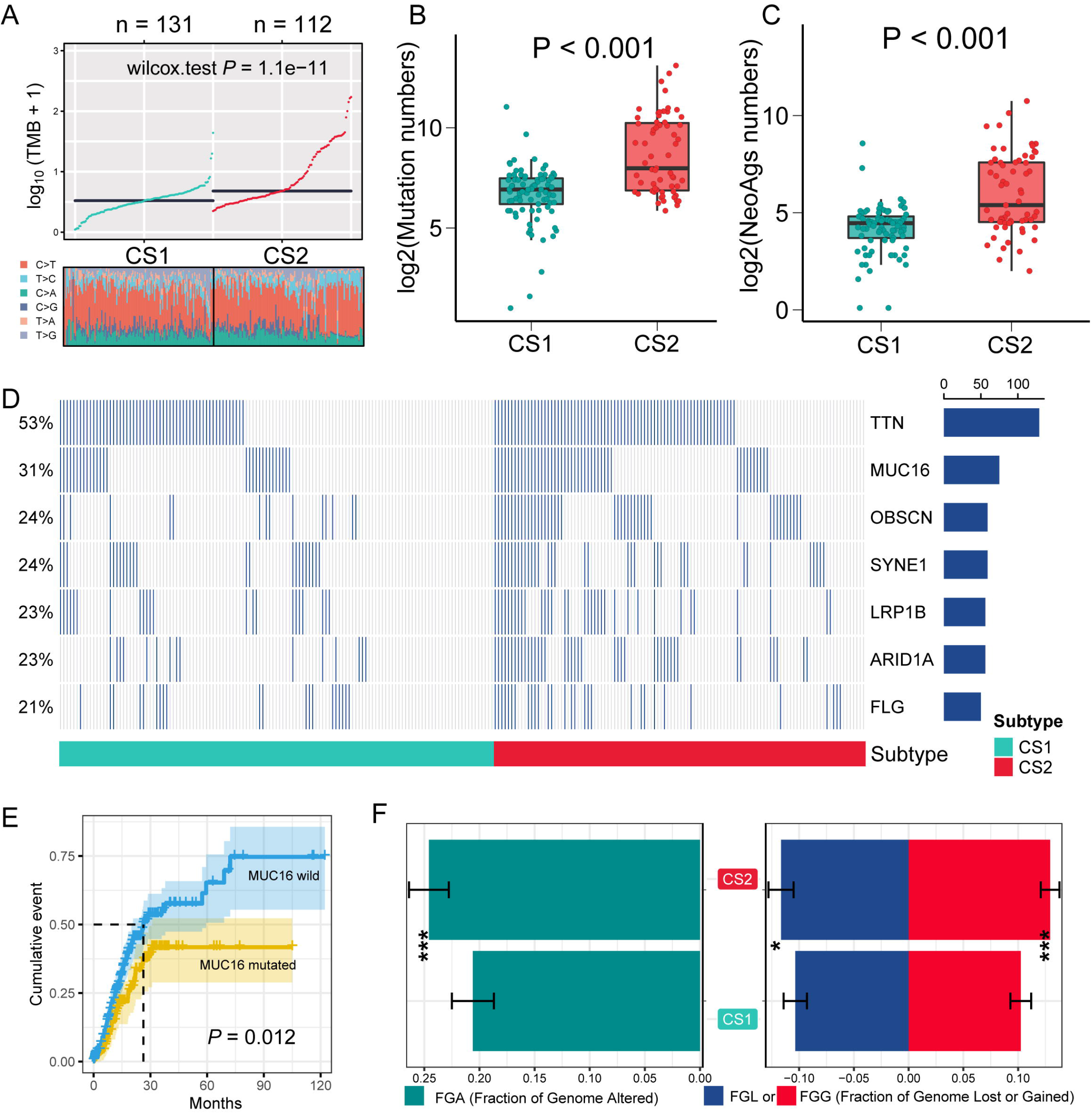
Diversely genetic alterations among two clusters. A. Comparison of TMB and TiTv among two identified groups of gastric cancer; B. The differential number of gene mutations in the CS1 and CS2 groups; C. The differential number of neoantigens in CS1 and CS2 groups; D. The landscape and top different mutant genes in the CS1 and CS2 groups; E. Cumulative events of death in patients with or without MUC16 mutation; F. Bar plot of fraction genome altered among two clusters.

### CS2 group contained more activated immune infiltration and responded more from immuno-/chemotherapy

Due to the CS2 group contained more gene mutations, CNAs and neo antigens. Hence, it is reasonable to postulate that the CS2 group underwent an activated status of immune infiltration. Based on the calculation of GSEA to immune activation and suppression signatures, the CS2 subtype was found to contain a higher score of activation signatures, including immune enrichment score, cytotoxic cells, activated CD4 T cell, activated CD8 T cell, 6 gene IFN signature, and CYT. Meanwhile, an activated status of suppression signatures in the CS1 group was also observed, including MDSC, Macrophages, Wnt/TGFβ signature, TITR, TGFB1 activated and C-ECM signatures (**Figure 6A**). Furthermore, the potential response to immunotherapy of these 243 GC patients were evaluated by TIDE, in which a total of 65 patients served as potential responders to immunotherapy (**Figure 6B**), while more responders belonged to the CS2 group (32% vs. 22%, P = 0.083, **Figure 6C**). However, the potential response to chemo drugs commonly used in GC patients was also assessed, which showed that the CS2 group responded more to treatment with 5-fluorouracil, cisplatin, and paclitaxel (all P < 0.01, **Figure 6D**).

**Figure 6.**
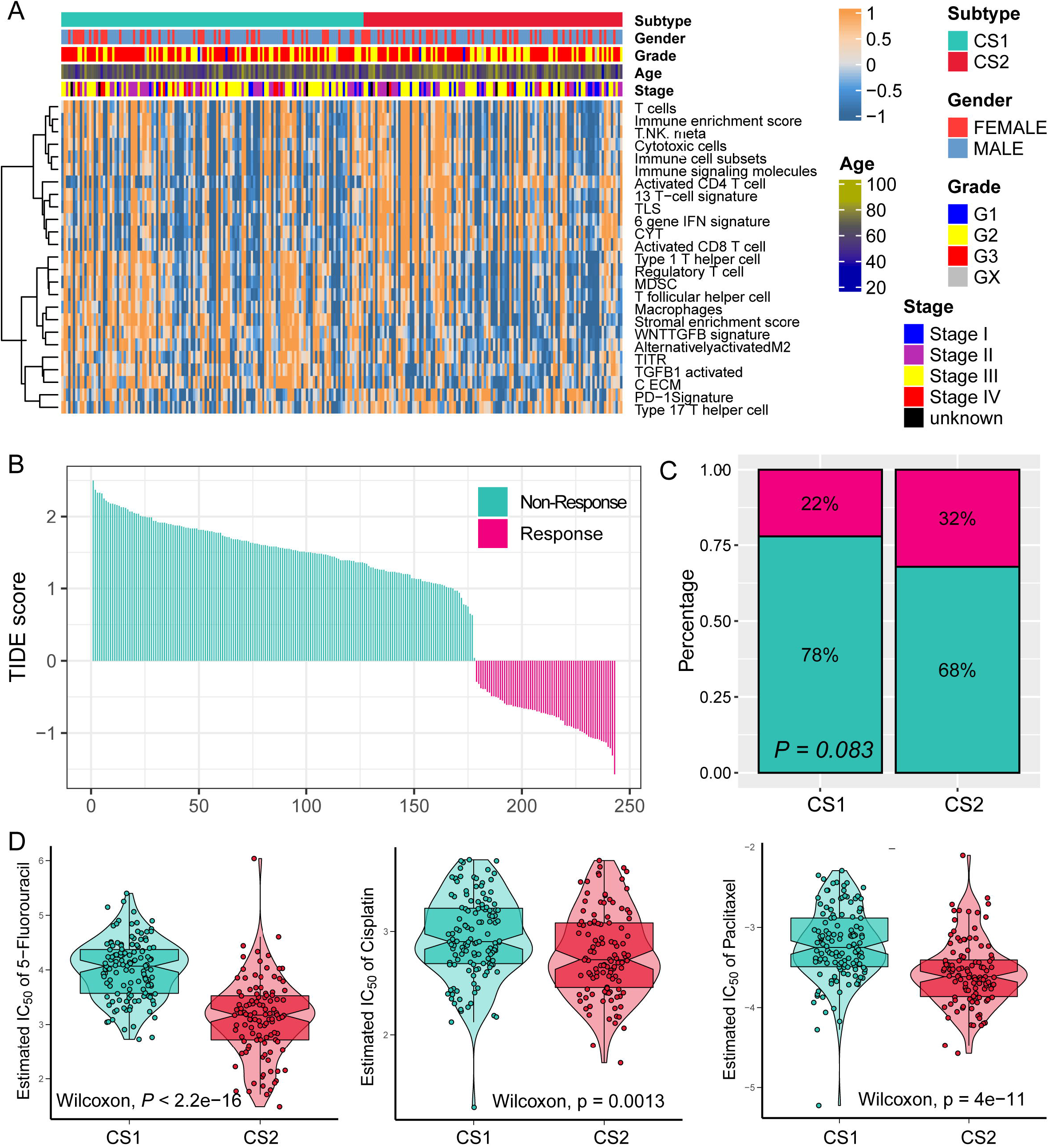
Potential treatment targets for gastric patients in two clusters. A. The activation of immune associated signatures in the CS1 and CS2 subtypes; B. The distribution of immunotherapy responders and non-responders predicted by the TIDE method; C. Patients in the CS2 cluster seem to be more suitable for immunotherapy; D. 5-Fluorouracial, cisplatin and paclitaxel seem to be more suitable for CS2 patients.

### The successful validation of newly identified subtypes in four external cohorts

Based on the “limma” package, the top 200 upregulated biomarkers were identified (**Figure S2**) for CS1 and CS2, respectively, with a significance threshold (adjusted P < 0.05) and no overlap with any biomarkers identified for other subtypes. The four external cohorts were noted to be GSE62254, GSE26253, GSE15459, and GSE84437, containing a total of 1357 GC patients. By employing the NTP method, the CS1 and CS2 group was predicted for each cohort by subtype-specific upregulated biomarkers, respectively (**Figure 7A**). Notably, the consistent prognosis results of CS1 and CS2 were illustrated in all of the four external GEO cohorts. Patients in the CS1 group had a more unfavorable OS outcome, while those in the CS2 group faced a favorable prognosis (GSE62254, HR = 0.53, 95% CI = 0.38-0.72, P < 0.001; GSE26253, HR = 0.71, 95% CI = 0.52-0.96, P = 0.027; GSE15459, HR = 0.61, 95% CI = 0.41-0.92, P = 0.018; GSE84437, HR = 0.59, 95% CI = 0.45-0.78, P < 0.001; **Figure 7B**).

**Figure 7.**
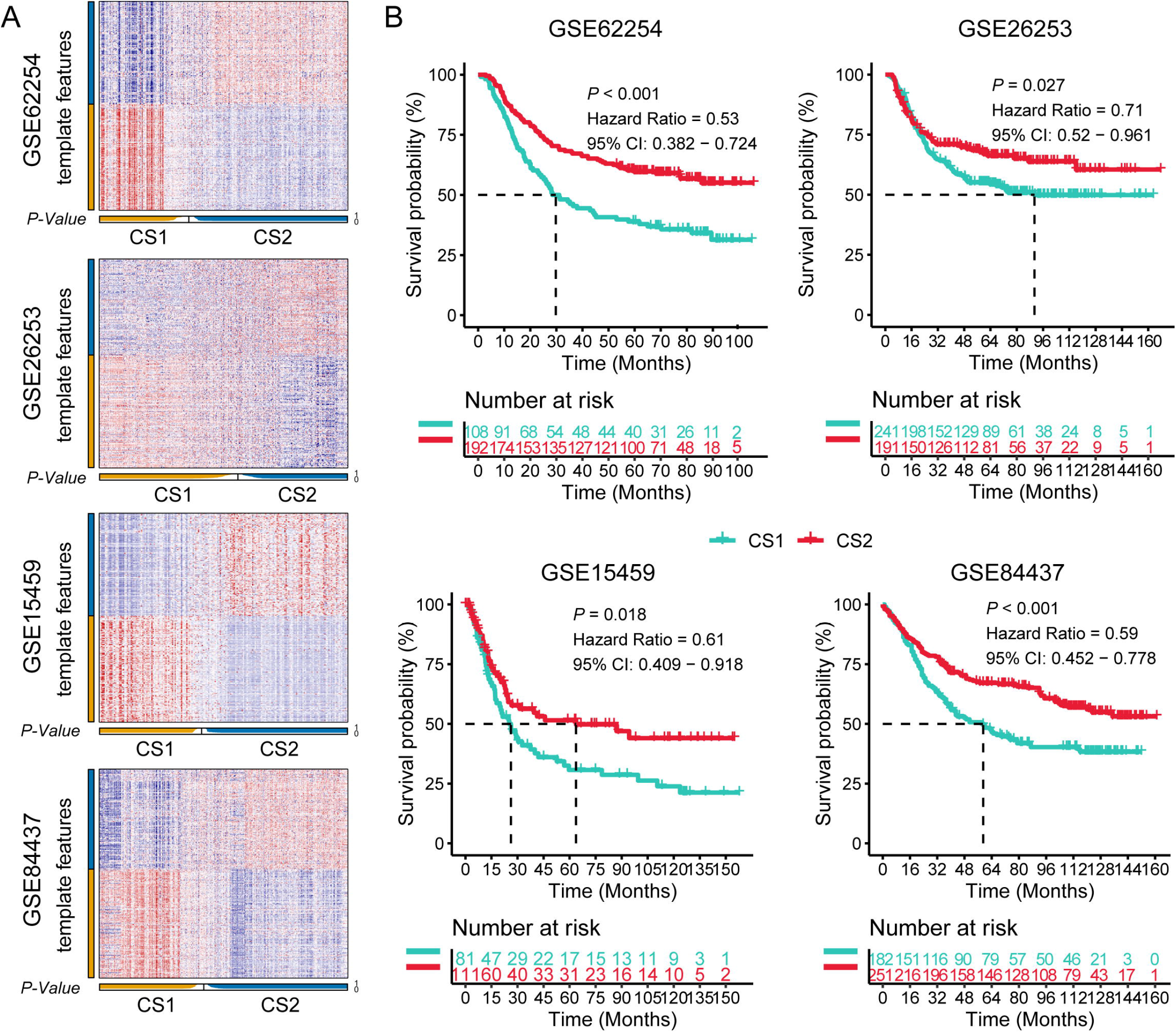
Successful validation of newly defined subtypes in four external cohorts. A. NTP method was applied to predict the CS1 and CS2 group for each cohort by the subtype-specific upregulated biomarkers; B. K-M plot showing the diverse overall survival in the CS1 and CS2 group of four external cohorts.

## Discussion

Several studies have reported the molecular subtypes of GC patients in order to illustrate the tumor heterogenies and guide clinical treatment. The Cancer Genome Atlas Research Network has proposed a molecular classification to divide gastric cancer into four subtypes: Epstein–Barr virus (EBV), microsatellite instability (MSI), genomically stable (GS) and chromosomal instability (CIN)[11]. EBV subtype displays recurrent PIK3CA mutations, extreme DNA hypermethylation, and amplification of JAK2, PD-L1 and PD-L2; MSI subtype shows elevated mutation rates; GS subtype is enriched for the diffuse histological variant and mutations of RHOA; and CIN subtype shows marked aneuploidy and focal amplification of receptor tyrosine kinases. Meng et al.[30] pointed out three GC subtypes that had different DNA methylation modification patterns based on the DNA methylation of 1619 GC patients, for which high DNA methylation score (DMS) was characterized by immune activation status, increased tumor mutation burden, and tumor neoantigens, which had a favorable prognosis. Meanwhile, the activation of the stroma and absence of immune cell infiltration were observed in the low DMS group, with relatively poor survival. Zhou et al. [31] also refined the molecular subtypes of GC patients by epigenetic age acceleration pattern. The definition of age acceleration (AA) is the difference between biological and chronological age. Patients with positive AA demonstrated better prognosis compared to those with negative AA. Furthermore, patients who belonged to the EBV subtype contained the highest average AA compared to MSI, GS and CIN subtypes. However, few studies were concerned with the classification of GC patients with multi-omics data.

In the current study, using the newly developed R package “MOVICS”, two molecular subtypes, CS1 and CS2, were identified by applying multi-omics data consistent of mRNA, LncRNA, miRNA, DNA methylation and gene mutation. The number of 2 clusters was selected by the prediction of CPI and Gaps-statistics analysis, and the subsequent separated from the total 243 patients were based on the consensus ensembles of 10 clustering algorithms. Interestingly, patients in the CS1 group had a poor prognosis in regard to their OS and RFS, while those in CS2 had a longer OS and RFS time. The activated extracellular matrix and epithelial mesenchymal transition pathways were observed in the CS1 group as well. The molecular pathogenesis of GC was also highly associated with E-cadherin, which harbor genetic and epigenetic abnormalities in both germline and sporadic GCs[32]. Xue et al.[33] put forward that the level of ZEB1 expression correlates with the degree of differentiation, metastasis to lymph nodes, and the stage of GC. In addition, suppression of ZEB1 may improve the expression of E-cadherin in GC cells and reduce the expression of vimentin. The correlation between SMOC2 and diseases has been widely reported. Jang et al.[34] reported that SMOC2 is a tumor suppressor in colorectal cancer and can elevate the expression of SMOC2 and suppress cell migration, colony and sphere formation. However, Lu et al.[35] reported on the suppression function in colon carcinoma invasion and migration of ARNTL2 via decreased SMOC2-EMT activity. In the current study, highly expressed SMOC2 in the poor prognosis CS1 group was also observed, which, on further study, confirmed that increased SMOC2 indicated poor prognosis in both TCGA-STAD and three GEO cohorts. Moreover, the expression of SMOC2 was closely associated with EMT markers.

The higher genetic alternations of the CS2 group were also observed more than that of the CS1 group. Specifically, there were more TTN mutations, MUC16 mutations and ARID1A mutations occurring in the CS2 favorable prognosis group. MUC16 serves as a type I transmembrane mucin protein that is comprised of 3 components: a C-terminal domain, a tandem repeat region, and an extracellular N-terminal section. Cancer antigen 125 (CA-125), which is used to monitor disease progression in ovarian cancer, is part of the tandem repeat domain[36]. MUC16 expression is associated with proliferation and metastatic potential in tumor cells, and MUC16 is known to inhibit the natural killer cell mediated lysis of tumors[37, 38]. The transcriptomic results from Li et al.’s study support the suggestion that MUC16 mutation may predict sensitivity to anti-PD-1 therapy; MUC16 mutant gastric tumors are immunologically “hot”[39]. Zhao et al. described that mutated MUC16 predicted a better prognosis in GC patients and was potentially associated with PI3K/Akt/mTOR signaling and Myc expression. Furthermore, they validated that the knockdown of MUC16 inhibited PI3K/Akt/mTOR signaling and reduced the protein level of Myc. Additionally, Yang et al.[40] reported the prognostic value of the combined mutations of MUC4, MUC16 and TTN, which rose to 0.782 in the TCGA-STAD cohort and 0.735 in the external validation FUSCC cohort. In the current study, more MUC16 mutations for good prognosis of the CS2 subtype were identified, in which mutated MUC16 were found to reduce death events for GC patients. Also, the CS2 group with activated infiltration of immunocytes, high expression of PD-1, and high response rate to immunotherapy was revealed.

There are several new findings and advantages of note in the current study. First, subtypes with diverse clinical outcome, immune infiltration status, genetic alterations were identified using integrative analysis of multi-omics data. Second, 10 clustering algorithms were enrolled, which further generated integration subtypes via the consensus ensembles. Therefore, the current classification is more robust than other studies, which only used one type of classification method. Third, it was revealed that the CS1 group had a poorer prognosis with less genetic alterations and activation of EMT associated pathways, whereas the CS2 group contained high TMB, more mutations and CNAs, though it had a favorable prognosis and high response rate to immunotherapy. Fourth, the diverse OS outcome between CS1 and CS2 subtypes were successfully validated in a total of 1357 GC patients from four cohorts, demonstrating the universal applicability of the new classification for clinical use.

## Conclusion

In this study, novel insight into the GC subtypes was obtained via integrative analysis of five omics data by utilizing ten state-of-the-art multi-omics clustering algorithms. Accordingly, patients in the CS1 group were found to have a poor prognosis and were linked with activation of EMT-associated signaling, while those in the CS2 group had a high mutation rate, high level of infiltration of activated immunocytes, and high response rate to immunotherapy.

## Acknowledgements

This work was supported by the National Natural Science Foundation of China (grant numbers: 81972539); the Key Research and Development Projects of Anhui Province (grant numbers: 201904a07020055).

## Author contributions

Conception and Design: Xianyu Hu, Zhenglin Wang, Juan Du and Wei Chen. Collection and Assembly of Data: Qing Wang, Ke Chen, Qijun Han and Suwen Bai. Data Analysis and Interpretation: Xianyu Hu, Zhenglin Wang, Ke Chen and Suwen Bai. Manuscript Writing: Xianyu Hu, Qing Wang, Qijun Han and Wei Chen. Final Approval of Manuscript: All the authors.

## Availability of data and materials

All data used in this work can be acquired from the GDC portal (https://portal.gdc.cancer.gov/), Gene-Expression Omnibus (GEO; https://www.ncbi.nlm.nih.gov/geo/).

## References

1. Bray F, Ferlay J, Soerjomataram I, Siegel RL, Torre LA, Jemal A: Global cancer statistics 2018: GLOBOCAN estimates of incidence and mortality worldwide for 36 cancers in 185 countries. CA Cancer J Clin 2018, 68(6):394–424.

2. Gao K, Wu J: National trend of gastric cancer mortality in China (2003-2015): a population-based study. Cancer Commun (Lond) 2019, 39(1):24.

3. Wang FH, Shen L, Li J, Zhou ZW, Liang H, Zhang XT, Tang L, Xin Y, Jin J, Zhang YJ et al: The Chinese Society of Clinical Oncology (CSCO): clinical guidelines for the diagnosis and treatment of gastric cancer. Cancer Commun (Lond) 2019, 39(1):10.

4. Hatakeyama M: Malignant Helicobacter pylori-Associated Diseases: Gastric Cancer and MALT Lymphoma. Adv Exp Med Biol 2019, 1149:135–149.

5. Ono H, Yao K, Fujishiro M, Oda I, Nimura S, Yahagi N, Iishi H, Oka M, Ajioka Y, Ichinose M et al: Guidelines for endoscopic submucosal dissection and endoscopic mucosal resection for early gastric cancer. Dig Endosc 2016, 28(1):3–15.

6. Das M: Neoadjuvant chemotherapy: survival benefit in gastric cancer. Lancet Oncol 2017, 18(6):e307.

7. Fujita K, Kanda M, Ito S, Mochizuki Y, Teramoto H, Ishigure K, Murai T, Asada T, Ishiyama A, Matsushita H et al: Association between Lymphovascular Invasion and Recurrence in Patients with pT1N+ or pT2-3N0 Gastric Cancer: a Multi-institutional Dataset Analysis. J Gastric Cancer 2020, 20(1):41–49.

8. Santoro R, Ettorre GM, Santoro E: Subtotal gastrectomy for gastric cancer. World J Gastroenterol 2014, 20(38):13667–13680.

9. Choi AH, Kim J, Chao J: Perioperative chemotherapy for resectable gastric cancer: MAGIC and beyond. World J Gastroenterol 2015, 21(24):7343–7348.

10. Ilson DH: Advances in the treatment of gastric cancer: 2019. Curr Opin Gastroenterol 2019, 35(6):551–554.

11. Cancer Genome Atlas Research N: Comprehensive molecular characterization of gastric adenocarcinoma. Nature 2014, 513(7517):202–209.

12. Xing X, Guo J, Ding G, Li B, Dong B, Feng Q, Li S, Zhang J, Ying X, Cheng X et al: Analysis of PD1, PDL1, PDL2 expression and T cells infiltration in 1014 gastric cancer patients. Oncoimmunology 2018, 7(3):e1356144.

13. Pietrantonio F, Miceli R, Raimondi A, Kim YW, Kang WK, Langley RE, Choi YY, Kim KM, Nankivell MG, Morano F et al: Individual Patient Data Meta-Analysis of the Value of Microsatellite Instability As a Biomarker in Gastric Cancer. J Clin Oncol 2019, 37(35):3392–3400.

14. Usui G, Matsusaka K, Mano Y, Urabe M, Funata S, Fukayama M, Ushiku T, Kaneda A: DNA Methylation and Genetic Aberrations in Gastric Cancer. Digestion 2021, 102(1):25–32.

15. Derks S, de Klerk LK, Xu X, Fleitas T, Liu KX, Liu Y, Dietlein F, Margolis C, Chiaravalli AM, Da Silva AC et al: Characterizing diversity in the tumor-immune microenvironment of distinct subclasses of gastroesophageal adenocarcinomas. Ann Oncol 2020, 31(8):1011–1020.

16. Cristescu R, Lee J, Nebozhyn M, Kim KM, Ting JC, Wong SS, Liu J, Yue YG, Wang J, Yu K et al: Molecular analysis of gastric cancer identifies subtypes associated with distinct clinical outcomes. Nat Med 2015, 21(5):449–456.

17. Fashoyin-Aje L, Donoghue M, Chen H, He K, Veeraraghavan J, Goldberg KB, Keegan P, McKee AE, Pazdur R: FDA Approval Summary: Pembrolizumab for Recurrent Locally Advanced or Metastatic Gastric or Gastroesophageal Junction Adenocarcinoma Expressing PD-L1. Oncologist 2019, 24(1):103–109.

18. Lordick F, Shitara K, Janjigian YY: New agents on the horizon in gastric cancer. Ann Oncol 2017, 28(8):1767–1775.

19. Pereira MA, Ramos M, Dias AR, Faraj SF, Ribeiro RRE, de Castria TB, Zilberstein B, Alves VAF, Ribeiro U, Jr., de Mello ES: Expression Profile of Markers for Targeted Therapy in Gastric Cancer Patients: HER-2, Microsatellite Instability and PD-L1. Mol Diagn Ther 2019, 23(6):761–771.

20. Rodriquenz MG, Roviello G, D’Angelo A, Lavacchi D, Roviello F, Polom K: MSI and EBV Positive Gastric Cancer’s Subgroups and Their Link With Novel Immunotherapy. J Clin Med 2020, 9(5).

21. Yan HHN, Siu HC, Law S, Ho SL, Yue SSK, Tsui WY, Chan D, Chan AS, Ma S, Lam KO et al: A Comprehensive Human Gastric Cancer Organoid Biobank Captures Tumor Subtype Heterogeneity and Enables Therapeutic Screening. Cell Stem Cell 2018, 23(6):882–897 e811.

22. Lu X, Meng J, Zhou Y, Jiang L, Yan F: MOVICS: an R package for multi-omics integration and visualization in cancer subtyping. Bioinformatics 2020.

23. Chalise P, Fridley BL: Integrative clustering of multi-level ‘omic data based on non-negative matrix factorization algorithm. PloS one 2017, 12(5):e0176278.

24. Hastie T, Tibshirani R, Walther G: Estimating the number of data clusters via the Gap statistic. J Roy Stat Soc B 2001, 63:411–423.

25. Meng J, Lu X, Zhou Y, Zhang M, Ge Q, Zhou J, Hao Z, Gao S, Yan F, Liang C: Tumor immune microenvironment-based classifications of bladder cancer for enhancing the response rate of immunotherapy. Mol Ther Oncolytics 2021, 20:410–421.

26. Chen PL, Roh W, Reuben A, Cooper ZA, Spencer CN, Prieto PA, Miller JP, Bassett RL, Gopalakrishnan V, Wani K et al: Analysis of Immune Signatures in Longitudinal Tumor Samples Yields Insight into Biomarkers of Response and Mechanisms of Resistance to Immune Checkpoint Blockade. Cancer discovery 2016, 6(8):827–837.

27. Hoshida Y, Brunet JP, Tamayo P, Golub TR, Mesirov JP: Subclass mapping: identifying common subtypes in independent disease data sets. PloS one 2007, 2(11):e1195.

28. Rooney MS, Shukla SA, Wu CJ, Getz G, Hacohen N: Molecular and genetic properties of tumors associated with local immune cytolytic activity. Cell 2015, 160(1-2):48–61.

29. Mayakonda A, Lin DC, Assenov Y, Plass C, Koeffler HP: Maftools: efficient and comprehensive analysis of somatic variants in cancer. Genome Res 2018, 28(11):1747–1756.

30. Meng Q, Lu YX, Ruan DY, Yu K, Chen YX, Xiao M, Wang Y, Liu ZX, Xu RH, Ju HQ et al: DNA methylation regulator-mediated modification patterns and tumor microenvironment characterization in gastric cancer. Mol Ther Nucleic Acids 2021, 24:695–710.

31. Zhou YJ, Lu XF, Meng JL, Wang QW, Chen JN, Zhang QW, Zheng KI, Rocha CS, Martins CB, Yan FR et al: Specific epigenetic age acceleration patterns among four molecular subtypes of gastric cancer and their prognostic value. Epigenomics 2021, 13(10):767–778.

32. Bure IV, Nemtsova MV, Zaletaev DV: Roles of E-cadherin and Noncoding RNAs in the Epithelial-mesenchymal Transition and Progression in Gastric Cancer. Int J Mol Sci 2019, 20(12).

33. Xue Y, Zhang L, Zhu Y, Ke X, Wang Q, Min H: Regulation of Proliferation and Epithelial-to-Mesenchymal Transition (EMT) of Gastric Cancer by ZEB1 via Modulating Wnt5a and Related Mechanisms. Med Sci Monit 2019, 25:1663–1670.

34. Jang BG, Kim HS, Bae JM, Kim WH, Kim HU, Kang GH: SMOC2, an intestinal stem cell marker, is an independent prognostic marker associated with better survival in colorectal cancers. Sci Rep 2020, 10(1):14591.

35. Lu M, Huang L, Tang Y, Sun T, Li J, Xiao S, Zheng X, Christopher O, Mao H: ARNTL2 knockdown suppressed the invasion and migration of colon carcinoma: decreased SMOC2-EMT expression through inactivation of PI3K/AKT pathway. Am J Transl Res 2020, 12(4):1293–1308.

36. Felder M, Kapur A, Gonzalez-Bosquet J, Horibata S, Heintz J, Albrecht R, Fass L, Kaur J, Hu K, Shojaei H et al: MUC16 (CA125): tumor biomarker to cancer therapy, a work in progress. Mol Cancer 2014, 13:129.

37. Gubbels JA, Felder M, Horibata S, Belisle JA, Kapur A, Holden H, Petrie S, Migneault M, Rancourt C, Connor JP et al: MUC16 provides immune protection by inhibiting synapse formation between NK and ovarian tumor cells. Mol Cancer 2010, 9:11.

38. Muniyan S, Haridas D, Chugh S, Rachagani S, Lakshmanan I, Gupta S, Seshacharyulu P, Smith LM, Ponnusamy MP, Batra SK: MUC16 contributes to the metastasis of pancreatic ductal adenocarcinoma through focal adhesion mediated signaling mechanism. Genes Cancer 2016, 7(3-4):110–124.

39. Li X, Pasche B, Zhang W, Chen K: Association of MUC16 Mutation With Tumor Mutation Load and Outcomes in Patients With Gastric Cancer. JAMA Oncol 2018, 4(12):1691–1698.

40. Zhao H, Zhang L: MUC16 mutation predicts a favorable clinical outcome and correlates decreased Warburg effect in gastric cancer. Biochem Biophys Res Commun 2018, 506(4):780–786.

